# IMMUNOTAR - Integrative prioritization of cell surface targets for cancer immunotherapy

**DOI:** 10.1101/2024.06.04.597422

**Authors:** Rawan Shraim, Brian Mooney, Karina L. Conkrite, Amber K. Weiner, Gregg B. Morin, Poul H. Sorensen, John M. Maris, Sharon J. Diskin, Ahmet Sacan

## Abstract

Cancer remains a leading cause of mortality globally. Recent improvements in survival have been facilitated by the development of less toxic immunotherapies; however, identifying targets for immunotherapies remains a challenge in the field. To address this challenge, we developed IMMUNOTAR, a computational tool that systematically prioritizes and identifies candidate immunotherapeutic targets. IMMUNOTAR integrates user-provided RNA-sequencing or proteomics data with quantitative features extracted from publicly available databases based on predefined optimal immunotherapeutic target criteria and quantitatively prioritizes potential surface protein targets. We demonstrate the utility and flexibility of IMMUNOTAR using three distinct datasets, validating its effectiveness in identifying both known and new potential immunotherapeutic targets within the analyzed cancer phenotypes. Overall, IMMUNOTAR enables the compilation of data from multiple sources into a unified platform, allowing users to simultaneously evaluate surface proteins across diverse criteria. By streamlining target identification, IMMUNOTAR empowers researchers to efficiently allocate resources and accelerate immunotherapy development.

## Introduction

In the last decade, more targeted and less toxic cancer therapies have gained prominence in cancer treatment [1]. Immunotherapy specifically has revolutionized cancer treatment by leveraging an individual’s immune system to combat cancer cells through natural mechanisms that are impeded during disease progression [2, 3]. Several types of immunotherapies have shown success in cancer treatments such as immune checkpoint inhibitors (ICIs), which have shown remarkable effectiveness in tumors with a high mutational burden such as melanoma and lung cancers; however, ICIs remain ineffective in most low mutational burden cancers such as childhood malignancies [4]. Immunotherapies that engage T-lymphocytes, including chimeric antigen receptor (CAR) T-cell therapy, on the other hand, have shown notable efficacy and durable responses in hematological cancers for both pediatric and adult patients and are beginning to show signs of efficacy in solid tumors [5–7]. Antibody drug conjugates (ADCs), a type of monoclonal antibody immunotherapy that delivers cytotoxic drugs directly to the tumor, have proven efficacy in hematological tumors and in solid tumors, with a few already gaining FDA approval and many more in the clinical development pipeline [7].

Development of successful immunotherapies such as CAR-T cell therapy and ADCs involve identifying cancer-specific proteins to target. These proteins are traditionally surface proteins that arise from the accumulation of genetic or epigenetic aberrations which lead to expression of proteins that are either not found on normal tissue, or are present at elevated levels on tumor cells and very low levels on healthy cells, or are only present on germ and tumor cells [6, 8, 9]. Technologies such as RNA-sequencing and mass spectrometry have been employed to explore the landscape of cell surface protein expression of cancers to identify these tumor-specific proteins [10–13].

Several criteria exist for an ideal surface protein immunotherapeutic target including high and homogeneous expression in cancer, minimal expression in normal healthy tissues, robust confidence in surface protein localization and functional relevance to the tumor [8, 14]. Protein glycosylation is another factor to consider when selecting targets, as glycosylation patterns can be altered in cancer cells, leading to the presentation of cancer-specific epitopes [15]. While publicly available databases allow investigation of these “ideal” surface protein characteristics, no tool currently enables users to query multiple databases simultaneously and systematically evaluate a protein’s suitability as an immunotherapeutic target.

To address the urgent need to identify surface protein targets for therapeutic development, we developed IMMUNOTAR, a tool designed to handle user-provided cancer RNA or proteomic expression datasets, query each cancer protein across multiple public databases that tackle some of the criteria for an “ideal” therapeutic protein, and produce a score that quantitatively evaluates each protein as a potential immunotherapeutic candidate.

## Results

### Overview of IMMUNOTAR

IMMUNOTAR is a tool developed in the R programming language and is designed to integrate user-provided proteomic or RNA-sequencing expression datasets with publicly available databases. The goal of IMMUNOTAR is to assign scores to proteins representing their predicted suitability as an immunotherapeutic target. We equipped IMMUNOTAR with both adult-derived and pediatric-specific databases to aid the development of immunotherapy in childhood cancers as well as adult cancers. The databases included within IMMUNOTAR are selected to tackle the ideal immunotherapeutic target criteria and can be divided into four categories: normal tissue expression, protein localization, biological annotation, and reagent/therapeutic availability (**Fig. 1A) (Supplemental Table S1)**. For normal tissue expression we included adult-derived RNA-sequencing data from the Genotype-Tissue Expression (GTEx) and pediatric-derived RNA-sequencing data from the Evo-Devo Mammalian organs (Evo-Devo) project [16, 17]. Acknowledging potential disparities between RNA expression and protein abundance, we also incorporated proteomics data derived from adult-normal human tissues generated by Jiang and colleagues [18]. For protein localization information, we included data from Compiled Interactive Resource for Extracellular and Surface Studies (CIRFESS), COMPARTMENTS, and UniProt [19–21]. To gain deeper insights into the biological roles of protein targets, data from the DepMap project and Gene Ontology (GO) were added [22, 23]. Identifying novel immunotherapeutic targets is inherently challenging due to the high possibility of poor or non-existent reagents, as highlighted in other studies [12]. To address this, we have incorporated data from the Therapeutic Target Database (TTD), the Database of Antibody-drug Conjugates (ADCdb), and the National Cancer Institute (NCI) relevant Pediatric Molecular Targets List (PMTL) https://moleculartargets.ccdi.cancer.gov/mtp-pmtl-docs), which helps users determine if the targets they are interested in have been identified and validated in other disease phenotypes [24, 25]. This integration facilitates the prioritization of targets with established reagents, enhancing the potential for successful validation and development.

**Figure 1:**
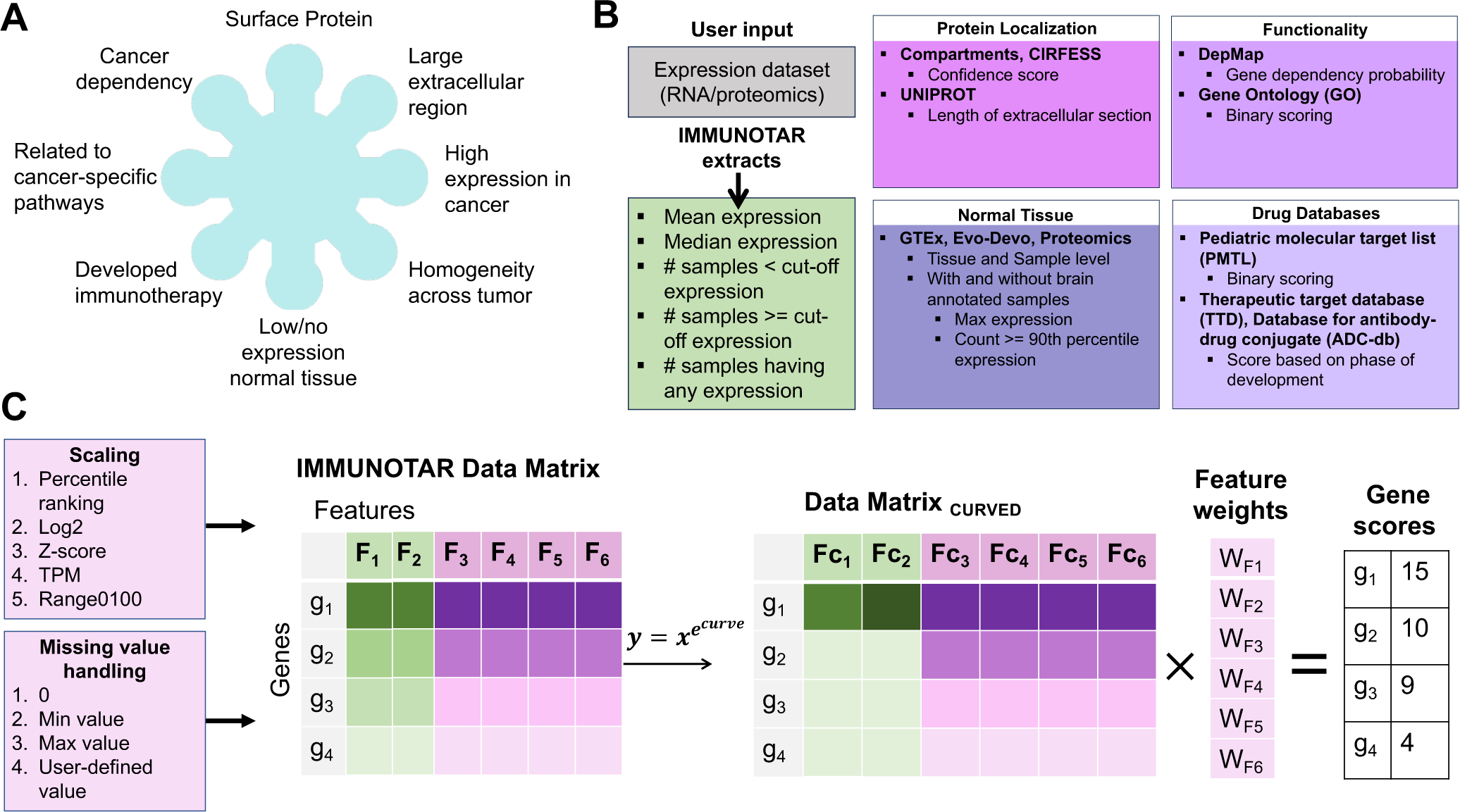
IMMUNOTAR feature extraction and surface protein scoring scheme. **A**) The pre-defined criteria for an ideal immunotherapeutic target. **B**) Summary of features extracted from the user-input expression dataset and the publicly available databases incorporated in IMMUNOTAR. **C**) The steps of analysis within IMMUNOTAR. The first step involves extracting quantitative features from the user input and the integrated databases to form a scoring data matrix. In the second step, IMMUNOTAR can rescale and handle missing values of the extracted features using the methods listed. After rescaling and handling missing values of the features, the user has the option of applying a curving function to the feature values. After curving the feature values, a weight parameter can be applied and the final score of each gene is calculated as the sum of the weighted-curved-features.

IMMUNOTAR analyzes a dataset (project) in 3 main steps: (1) Generating a gene-by-features data matrix, (2) Applying project analysis parameters to the data-matrix, and (3) Calculating the IMMUNOTAR gene score and evaluating how the algorithm performed in prioritizing targets. Project analysis parameters are customizable as IMMUNOTAR is designed to accept user-input for the parameters using a project parameters file (**Supplemental File S1)**.

Step one of the analyses involves extracting summary features for each gene in the expression dataset that is provided by the user and enriching with quantitative features from the public databases that are incorporated within IMMUNOTAR (**Fig. 1B)** (**Supplemental Table S2**). The output from step one is a genes-by-features data-matrix (**Fig. 1C**). The user has the flexibility to select the summarization and enrichment database features that they desire to include in the genes-by-features data-matrix within the project analysis parameters file, with the default being to include all.

In step two, IMMUNOTAR applies additional project analysis parameters to the genes-by-features data-matrix. Initially, IMMUNOTAR will rescale and fill in any missing values for all features in the data-matrix (**Fig. 1C**). Next, IMMUNOTAR applies a curving parameter as a form of a non-linear normalization of the feature values. Feature-specific weights are then applied to produce the final feature value. The user can customize the parameter values associated with each of the features using the project analysis parameters file. Assigning different weights, curving, and scaling of feature values ensures that some features contribute more significantly to the final score of the gene in comparison to others.

The last step in any analysis is calculating the score for each gene using the weighted average of the final feature values (**Fig. 1C**). One of the outputs from IMMUNOTAR is a table that includes the feature values and the final score assigned to each gene. To evaluate IMMUNOTAR’s performance in prioritizing targets, the mean average precision score (MAP) is calculated and given as an output with the analysis (**Fig. 2A)**. The MAP score indicates how effectively immunotherapy targets in the queried cancer phenotype that are either in-clinic or in-development (known-positive targets) are ranked within IMMUNOTAR. The user can designate the known-positive targets in the project analysis parameters file or alternatively, they can be extracted using the therapeutic drug databases incorporated within IMMUNOTAR. If the user extracts this information using the available drug-databases, targets that have drugs that have been discontinued will be considered “known-negatives” and the MAP score will be calculated with those targets in consideration (**Fig. 2A)**.

**Figure 2:**
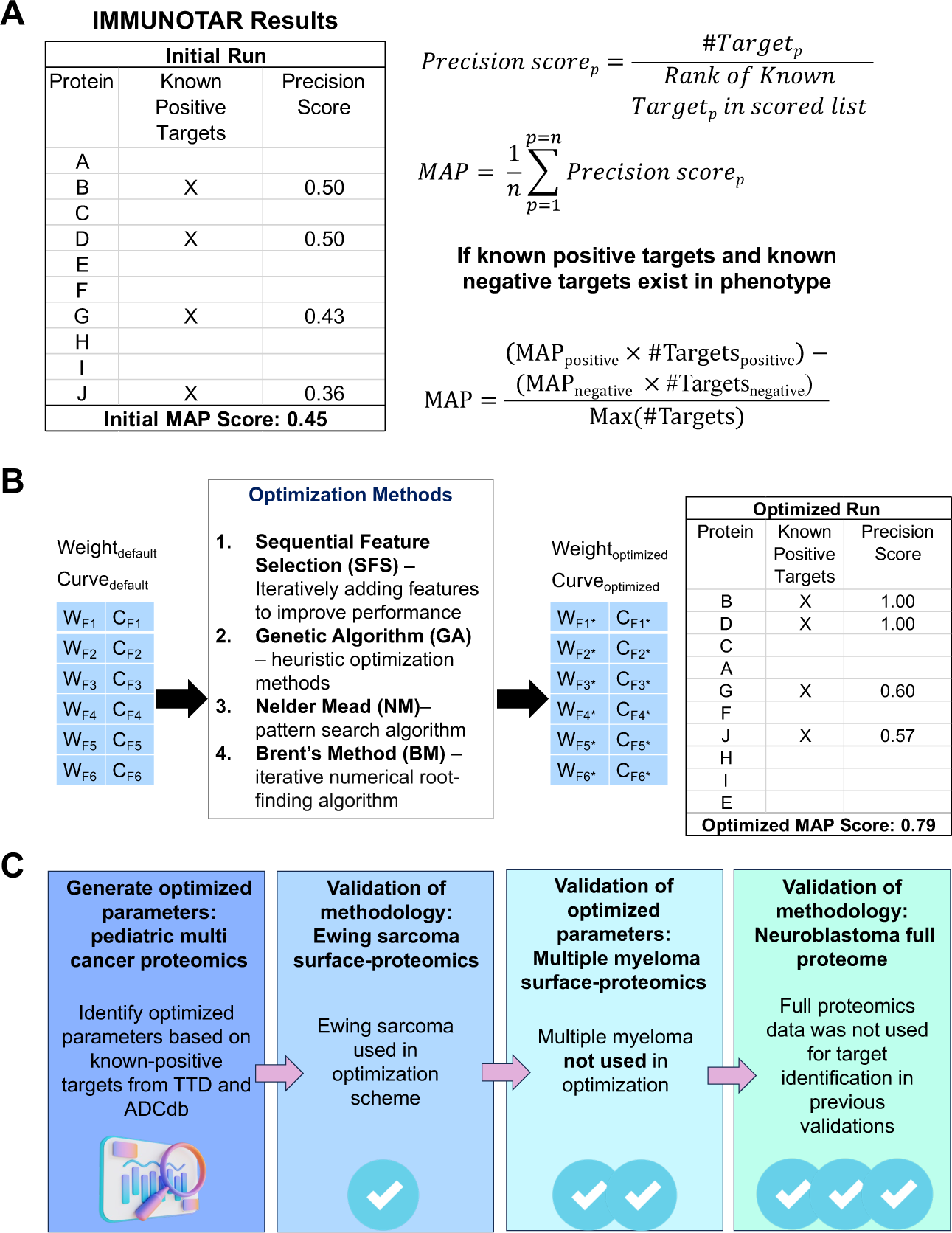
IMMUNOTAR evaluation, optimization, and validation datasets. **A)** The mean average precision (MAP) score is used to evaluate the performance of IMMUNOTAR. The MAP score depends on the known-positive and known-negative targets assigned to each phenotype and is calculated based on the ranking of those targets after running IMMUNOTAR. **B)** Optimization methods listed can be used to change the weights and curves of features. The goal of the optimization is to rank the known-positive targets higher in the results thus increasing the overall MAP score of the algorithm. **C)** Optimized parameters are generated using a multi-cancer proteomic dataset and applied to other datasets to validate the approach. Validation datasets include Ewing sarcoma surface proteomics, multiple myeloma surface proteomics, and neuroblastoma full proteome.

The curve and weight values influence the final feature values and thus the gene score and the MAP score of the algorithm. The weight and curving values can be assigned by users based on prior knowledge, though this can be somewhat subjective. To enhance the objectivity of these assignments, we have incorporated optimization methods within IMMUNOTAR, including supervised search, stochastic search, and non-linear optimization techniques (**Fig. 2B)**. The goal of implementing these optimization methods is to enhance the MAP score of IMMUNOTAR, allowing the identification of weight and curve parameter values that rank the known-positive targets at the top of prioritized lists.

### Validation and assessment approach

Through this work, we assessed IMMUNOTAR using diverse cancer datasets (**Fig. 2C**). We systematically generated optimized analysis parameter values using a full proteome dataset that surveyed twelve pediatric cancer phenotypes with known-positive targets. During this process, we assessed the analysis parameter values by comparing the MAP scores of the algorithm when using the default versus the optimized parameter values. Next, we validated IMMUNOTAR’s methodology by identifying potential targets across three different cancer phenotypes and datasets: surface proteomics for Ewing Sarcoma (EwS), surface proteomics for Multiple Myeloma (MM), and full proteomics for neuroblastoma (NBL) [11, 12, 26].

To evaluate IMMUNOTAR’s ability to identify relevant immunotherapeutic targets, we compared the prioritized targets within IMMUNOTAR to those identified by other scoring workflows in EwS and MM. This comparison involved ranking assessments and performing Gene-Set Enrichment Analysis (GSEA), testing the effectiveness of IMMUNOTAR in target selection and benchmarking it against existing strategies in the field. Finally, we analyzed the top targets identified by IMMUNOTAR in EwS, MM, and NBL datasets to confirm its potential in pinpointing viable immunotherapeutic targets, demonstrating its utility across different cancer types and data sources.

### Optimization using a pediatric multi-cancer dataset increases IMMUNOTAR’s MAP score across cancer phenotypes

The curving and the weight parameter values are crucial in calculating the final gene score in IMMUNOTAR. To identify optimized curve and weight values per feature, we utilized proteomic data from cell lines representing twelve pediatric cancer phenotypes with known-positive targets [18] (**Fig. 3A)**. We first applied the default project analysis parameters to each of the phenotypes (**Supplemental File S2)**. Known-positive cell surface therapeutic targets were extracted from TTD and ADCdb (**Supplemental Table S3**).

**Figure 3:**
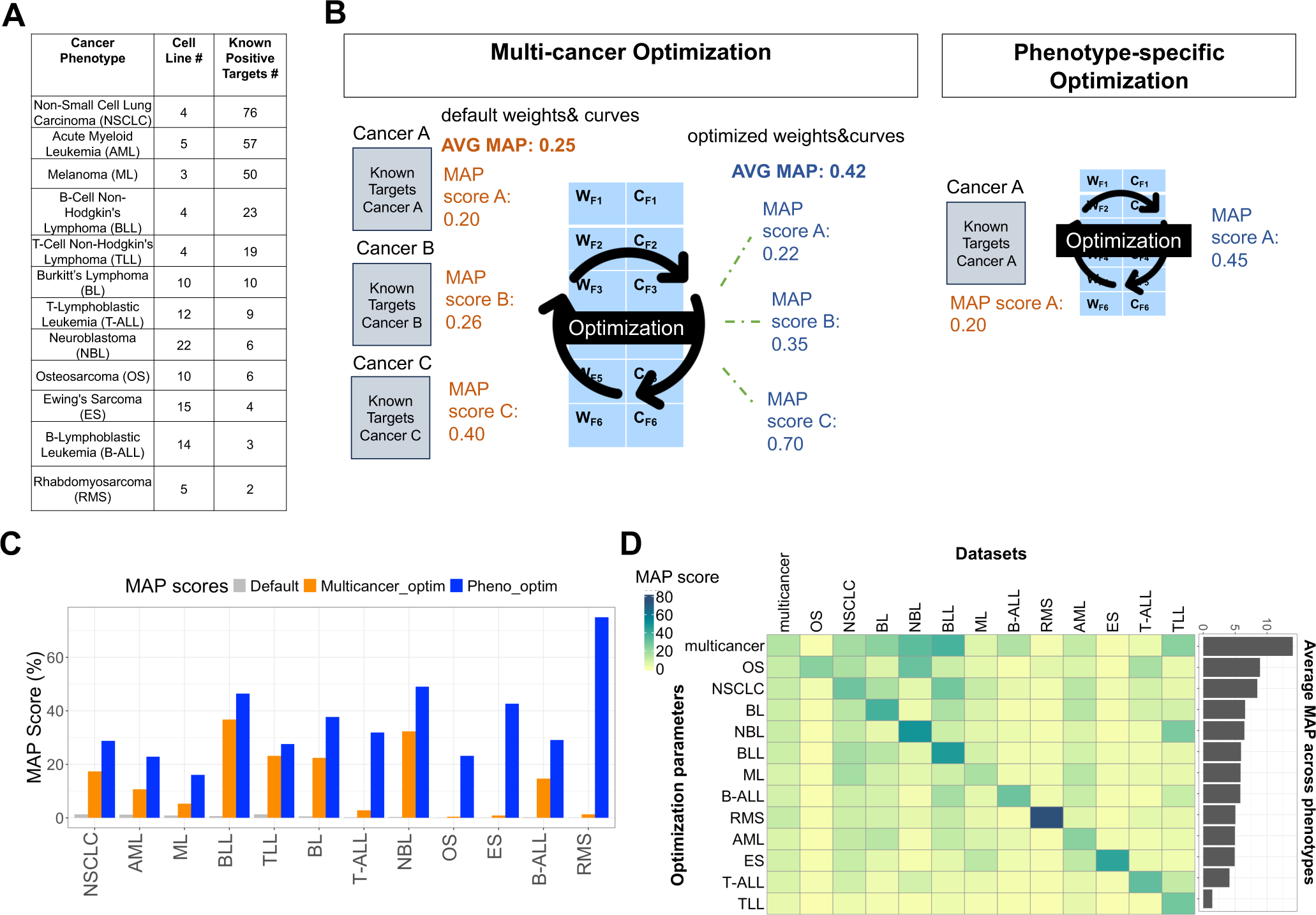
IMMUNOTAR optimization using pediatric pan-cancer expression data. **A)** The pediatric cancer phenotypes included in the optimization scheme, listing the number of cell lines and the number of known-positive surface protein targets extracted from the drug databases TTD and ADCdb (Supplemental Table S2). **B)** Optimization parameters were generated using two approaches, the first being a multi-cancer optimization approach and the second being a phenotype-specific optimization approach. **C)** Using the multi-cancer optimization parameters, the MAP score for each individual phenotype increased at varying levels within phenotypes. The phenotype-specific optimization approach increased the MAP score further for each phenotype. **D)** While the phenotype-specific parameters performed better on the phenotype itself, they did not perform as well on other phenotypes. Overall, the multi-cancer optimization parameters performed best across all phenotypes with an average MAP score of 14%.

We implemented two optimization strategies, including a multi-cancer optimization approach, where weights and curves values are evaluated across multiple cancer phenotypes at the same time and a phenotype-specific optimization approach, where weight and curve feature values are optimized using a single phenotype **(Fig. 3B).** To identify features contributing positively to the MAP score, we applied sequential forward selection combined with Brent’s optimization, a feature was included only if it improved the MAP score by 0.1%, and after inclusion, a search for the local maximum using Brent’s method was conducted [27]. The resulting optimization parameter values were then further refined using the genetic algorithm method, with a sub-optimization of the top-scoring parameters in the population using Nelder-Mead, only updating the weight or curve parameter values if it increased the MAP score by at least 0.1% [28, 29].

In the multi-cancer optimization strategy, only nine of the forty total features were retained by the feature selection method to calculate the final gene score (**Supplemental File S3**). After multi-cancer optimization, the average MAP score across all twelve phenotypes increased significantly by a 27-fold change (p-value < 0.001) in comparison to the default parameters, with an increase in the MAP score observed for every phenotype at varying degrees (**Fig. 3C)**. B cell non-Hodgkin’s Lymphoma and neuroblastoma exhibited the most significant increase in their MAP value post-optimization, reaching precision scores of 37% and 32%, respectively.

Our phenotype-specific optimization strategy increased the MAP score within each phenotype even further in comparison to the multi-cancer optimization strategy (**Fig. 3C)**. However, the phenotype-specific optimization parameters did not perform as well as the multi-cancer optimization parameters when applied to other phenotypes, showing that the average MAP score across phenotypes was the highest (14%) using the multi-cancer optimization parameter values (**Fig. 3D)**.

### IMMUNOTAR validates known EwS immunotherapy targets and highlights CADM1 as a new potential target

To test our optimization methodology and tool, we analyzed an EwS surface proteomics dataset from nineteen pediatric EwS patient-derived or cell line-derived xenograft models [12]. In the original publication, Mooney and colleagues filtered their data to 218 proteins within the generated surface and global proteome datasets based on protein localization criteria [12]. They prioritized and scored this subset using a custom, in-house workflow that is not fully automated, assigning a ranking score to their data and three public databases. Through their system they identified and validated novel targets including Ectonucleotide pyrophosphatase/-phosphodiesterase 1 (ENPP1) and Cadherin 11 (CDH11) [12].

Using IMMUNOTAR, phenotype-specific optimized weight and curve values were generated with the cell surface proteins prioritized by Mooney and colleagues as known-positive targets. The multi-cancer and phenotype-specific optimization parameter values were each applied and IMMUNOTAR results from both were compared. Using the multi-cancer optimization approach, ENPP1 and CDH11, validated novel targets by Mooney and colleagues, scored in the top 2% of targets (**Fig. 4A)** and using the phenotype-specific prioritization parameters, IL1RAP, a previously validated target within EwS, ENPP1 and CDH11 scored as the top three targets (**Fig. 4B**) (**Supplemental Tables S4-S5**). To compare scoring approaches, we completed a GSEA using three different subsets of targets (**Supplemental Table S6**). Using both the multi-cancer and phenotype-specific optimization parameters, IMMUNOTAR provided a significantly high enrichment score of 0.86 and 0.88 (p < 0.0001) for the prioritized targets, respectively (**Fig. 4C)**.

**Figure 4:**
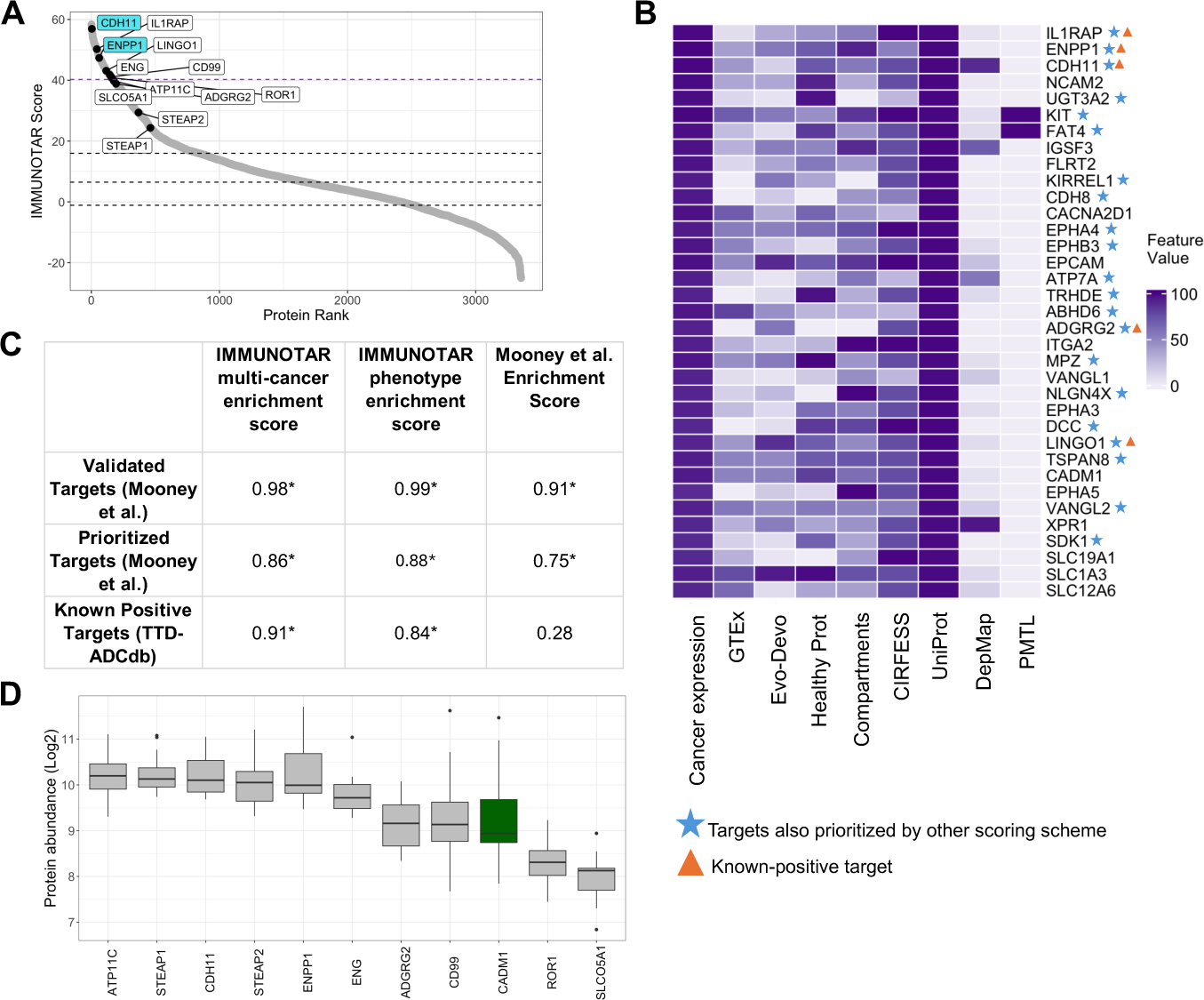
Validation of IMMUNOTAR using EwS surface proteomics data. **A)** IMMUNOTAR results from the EwS surface proteomics data using the multi-cancer optimization parameter values. The labeled proteins include targets that are known within EwS and highlighted by Mooney and colleagues. The blue labeled targets are novel targets discovered by Mooney and colleagues. The purple dashed line represents the cut-off for the top 5% of proteins scored by IMMUNOTAR. **B)** Heatmap showing the top 5% scored IMMUNOTAR targets after implementing a restricted normal-tissue expression filter within IMMUNOTAR. We marked targets that were prioritized by Mooney and colleagues scoring scheme and targets that were fed as known-positives in the IMMUNOTAR analysis. **C)** The GSEA enrichment scores comparing three sets of prioritized targets in EwS. The IMMUNOTAR scores of each target were compared to the target scores from the Mooney et al. scoring scheme. IMMUNOTAR outputs using both the multi-cancer optimization parameter values and the phenotype-specific optimization parameter values were compared to the Mooney et al. scoring scheme, separately. *p-value <= 0.05. **D)** Protein quantification of CADM1 in EwS surface proteomics dataset showing comparable abundance to other EwS-specific known-positive targets.

Filtering the output to highlight surface proteins with restricted normal tissue expression, defined as less than the 20^th^ percentile expression across all tissues measured by GTEx, IMMUNOTAR identified twenty-one targets that were also prioritized by Mooney and colleagues in the top 5^th^ scoring percentile (**Fig. 4B)** (**Supplemental Table S7**). Notably, nine targets were identified within IMMUNOTAR that scored in the top scoring bracket, z-score > 1, in the Mooney *et al.* scoring scheme including netrin receptor DDC (DCC), cadherin family member 14 (FAT4) and UDP-glucuronosyltransferase-3A2 (UGT3A2) (**Fig. 4B**).

A high scoring surface protein uniquely highlighted by IMMUNOTAR included cell adhesion molecule 1 (CADM1). CADM1 is a member of an immunoglobulin superfamily and has been involved in various tumor types such as ovarian cancer, breast cancer and osteosarcoma [30, 31]. Protein expression of CADM1 in the surface-proteomics dataset is comparable to other know-positive targets highlighted by Mooney and colleagues in their work (**Fig. 4D**). Although CADM1 is expressed in some normal tissues (**Supplemental Fig. S1**), studies in osteosarcoma have reported minimal toxicity with a CADM1-targeted ADC in *in-vivo* studies [30]. In small-cell lung cancer, researchers have identified a single-chain variable fragment specific to CADM1 in this phenotype and have developed an antibody therapy targeting it [31].

### IMMUNOTAR validates ITGA4, ITGB7, and FLVCR1 as candidate immunotherapeutic targets in MM

To verify the effectiveness of our multi-cancer optimization analysis parameters on phenotypes that were not initially included in IMMUNOTAR’s optimization process, we utilized a surface proteomics dataset from MM cell lines developed by Ferguson and colleagues [11]. Within their work, the group developed a manual ranking system where they subjectively assigned points to each criterion extracted from their experiment data and four public databases to score their candidates. The approach identified C-C motif Chemokine Receptor 10 (CCR10) as a novel candidate immunotherapeutic target [11]. Preliminary testing of this target showed that CCR10 could be a promising target for MM with some healthy tissue expression limitations [11]. They also highly ranked a second target, Thioredoxin Domain Containing 11 (TXNDC11), noting that MM cells depend on TXNDC11 for proliferation in DepMap [22]. Their validation of TXNDC11 revealed that the protein was highly expressed in MM cell lines; however, it showed primarily intracellular localization and was considered a false-positive of their scoring system [11].

Analyzing the MM surface proteomics data with IMMUNOTAR using the multi-cancer optimization parameters, CCR10 scored in the top 20 targets while TXNDC11 scored in the lower score quantiles, consistent with it representing a false positive in Ferguson *et al.* (**Fig. 5A) (Supplemental Table S8)**. Completing a GSEA showed that IMMUNOTAR had a higher enrichment score in comparison to their scoring algorithm when looking at known-positive targets from TTD and ADCdb (**Fig. 5B)**. Analyzing the proteins that scored in the top 2nd percentile, 13 out of 29 (44%) were identified in the literature as being relevant to MM, either as biomarkers of the disease, involved in pathways implicated in MM, or related to disease risk (**Fig. 5C)**. IMMUNOTAR ranked ITGA4 as the top scoring protein. ITGA4 has previously been implicated in myeloma cell homing, survival, and acquisition of cell-adhesion mediated drug resistance [32–35]. Another target within the top-ten was Integrin Subunit Beta7 (ITGB7), also known to enhance MM cell adhesion to bone-marrow stroma, migration, and invasion [36]. Clinically, expression of *ITGB7* in MM is associated with poor survival outcomes post autologous stem cell transplantation [36]. The GTEx expression of ITGB7 is limited to expression in the blood (**Fig. 4D)**. Feline Leukemia virus subgroup C receptor 1, FLVCR1, also in the top-ten scoring proteins, is a membrane heme exporter protein that has been reported to be possibly responsible for the poor prognosis associated with 1q32 gain in MM patients [37].

**Figure 5:**
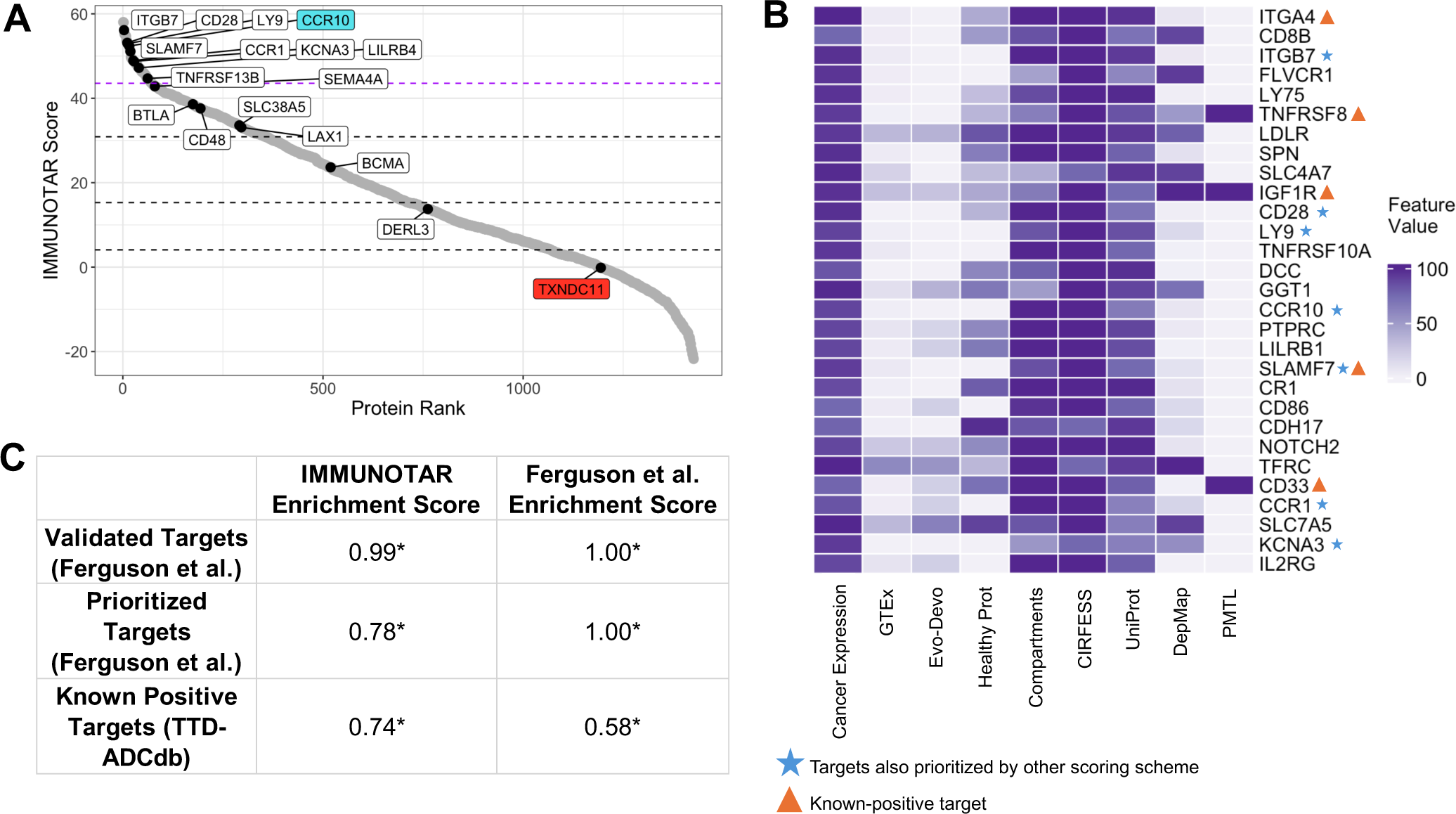
Validation of IMMUNOTAR using MM surface-proteomics data. **A)** IMMUNOTAR results from the MM surface proteomics dataset using the multi-cancer optimization parameters. Labeled proteins are ones that were ranked in the top-three scoring brackets by Ferguson and colleagues. The purple dashed line represents the top 5% scoring targets. The blue labeled target was prioritized and validated by Ferguson *et al.* as a novel target for MM. The red labeled target, TXNDC11 is a prioritized protein that was a false-positive proved by functional validation. **B)** The top 5% scoring targets using IMMUNOTAR, marking targets prioritized in IMMUNOTAR and the top-three scoring brackets and targets that are known-positive targets per TTD and ADCdb. **C)** The GSEA enrichment scores comparing three sets of prioritized targets in MM. The IMMUNOTAR scores of each target were compared to the target scores from the Ferguson et al. scoring scheme. *p-value <= 0.05.

### Phenotype-specific parameters validate known NBL immunotherapy targets and identify KCNH1 as novel candidate target

Finally, leveraging our expertise in NBL and utilizing a non-surface specific proteomics dataset, we applied IMMUNOTAR to cell-line based full proteome data that surveyed twenty-two NBL cell lines [26].

We generated phenotype-specific optimization parameters using known-positive targets that have been explored in the literature or within our group including ALK, GPC2, CD267, L1CAM, NCAM1, and DLK1 [13, 38–43]. Of note, DLK1 was not quantified in this NBL full proteome dataset and therefore was not evaluated. The multi-cancer optimization parameters achieved a MAP score of 32% (**Supplemental Table S9**). However, utilizing the phenotype-specific parameters improved the MAP score to 49%, ranking all known positives targets within the top 10^th^ percentile of proteins scored (**Supplemental Table S10**).

To identify new candidate immunotherapy targets in NBL, we focused on proteins with restricted expression in normal tissues, defined as less than 20th-percentile expression across all tissues measured by GTEx, leading to the identification of Potassium Voltage-Gated Channel Subfamily H Member 1 (KCNH1) (**Fig. 6A**) (**Supplemental Table S11**). Protein abundance of KCNH1 was comparable to targets under development in neuroblastoma including NCAM1, L1CAM and CD276, and higher than other known immunotherapeutic targets such as ALK and GPC2 (**Fig. 6B**). KCNH1 encodes the Kv10.1 potassium channel and is predominantly expressed in the brain but not in other normal tissues, as per GTEx data and Evo-devo dataset (**Fig. 6C**) (**Supplemental Fig. S2**). This restricted expression profile makes KCNH1 a good candidate for ADC development given that ADCs do not cross the blood-brain barrier, potentially avoiding off-target cytotoxic effects. KCNH1 overexpression has been implicated in cancer cell proliferation and tumor growth in cervical carcinoma and other soft tissue sarcomas [44, 45].

**Figure 6:**
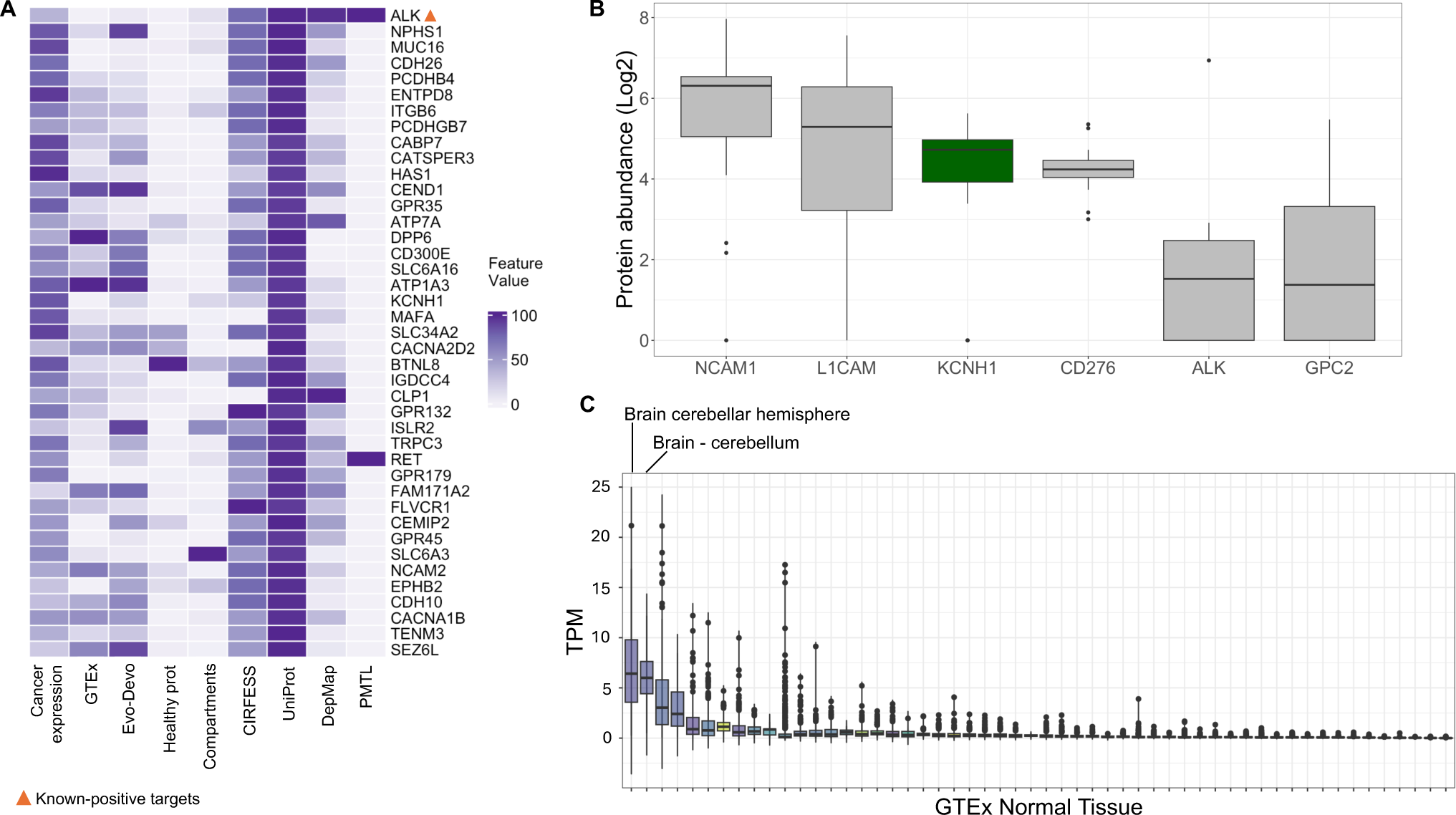
IMMUNOTAR prioritized targets in NBL using full proteome data and phenotype-specific optimization parameters. **A)** Evaluating the top 5% scoring targets per IMMUNOTAR analysis after applying phenotype-specific parameters and the restrictive normal-tissue expression filter. Highlighting known, top-scoring target, ALK. **B)** Protein quantification of KCNH1 in full proteome dataset showing comparable abundance to other NBL-specific known-targets. **C)** Normal tissue expression per GTEx for KCNH1 reveals limited normal tissue expression, with some expression in healthy brain. Tissues with median expression ≥ 5 TPM are labeled.

## Discussion

We present IMMUNOTAR, a tool designed to identify potential cell surface proteins for cancer immunotherapy development. IMMUNOTAR analyzes user-provided cancer RNA and proteomic datasets, integrating these with generated quantitative features from several public databases including, GTEx, Evo-Devo, CIRFESS, COMPARTMENTS, UniProt, DepMap, and GO. This integration allows users to perform comprehensive quantitative evaluation of potential targets without individually accessing each database. Additionally, IMMUNOTAR includes drug databases like TTD, ADCdb, and PMTL, enabling users to identify existing therapies and reagents for the prioritized targets. This utility facilitates validation and translational research, highlighting IMMUNOTAR’s bench-to-bedside potential.

To provide optimized analysis parameter values for users, we used a multi-cancer cell-line proteomics dataset and implemented both multi-cancer and phenotype-specific optimization strategies. Our analysis revealed that while phenotype-specific parameters performed best on their respective phenotypes, multi-cancer parameters offered broader applicability across different phenotypes. This suggests that multi-cancer optimization parameters are reliable for initial analyses, especially for cancers with few or no known positive cell-surface protein targets.

IMMUNOTAR was benchmarked against other scoring approaches which were effective but were not presented as tools available for use by the research community and lacked full automation and usability across various datasets. IMMUNOTAR successfully replicated the results of other groups. In MM, a cancer phenotype that had not been utilized to generate the optimized analysis parameters, IMMUNOTAR identified targets previously reported by other groups, confirming the applicability of our methodology and multi-cancer parameter values. Additionally, IMMUNOTAR identified novel potential targets in EwS and NBL, showcasing its ability to discover potential new therapeutic avenues.

Despite these successes, IMMUNOTAR relies on user-provided RNA or proteomics datasets, which can be affected by data sparsity or contamination, impacting protein quantification and target prioritization. For example, DLK1 was not quantified in the full proteome data for NBL but was identified in other surface-protein datasets [13]. Additionally, in some analyses, researchers are confined to using cell line models, due to limited patient sample availability, which can misrepresent true protein tumor expression [46]. These discrepancies underscore the importance of high-quality, comprehensive datasets for accurate target identification using IMMUNOTAR, as well as the necessity of validating findings on human samples.

Although our optimization strategy provides a standardized method for determining feature weights and curves, it has its limitations. Some targets within the TTD and ADCdb are still undergoing development and may lack finalized efficacy and safety results, potentially leading to the prioritization of false positives. This could result in the inappropriate assignment of weight and curve values to features that do not truly reflect an “ideal” target. Conversely, ranking discontinued targets as “known-negative” may not always accurately represent their potential, as therapy failure could be due to factors unrelated to the target itself.

In our analyses, we focus on using IMMUNOTAR to analyze surface proteomic data, as it provides a targeted quantification of surface proteins compared to RNA-sequencing and full-proteome data. Another promising technique is immunopeptidomics, a targeted mass-spectrometry approach that discovers noncanonical antigens, tumor antigens not derived from traditional protein-coding regions of the genome. These antigens can be tumor-specific and shared among patients, making them valuable for immunotherapy [47, 48]. As large-scale tumor and normal tissue immunopeptidomics become available, IMMUNOTAR could be extended to incorporate these datasets. Additionally, IMMUNOTAR could be extended to analyze short-read and long-read RNA-sequencing data for tumor-specific splice isoforms, as well as mass-spectrometry data for tumor-specific glycosylated proteins [49–51]. These enhancements would broaden the range of target selection, significantly improving IMMUNOTAR’s ability to identify cancer-specific and novel immunotherapeutic targets.

To increase the efficacy of immunotherapies and prevent cancer relapse and therapy resistance, researchers have explored designing immunotherapies that target multiple cancer-specific proteins (dual therapies) [52–54]. Dual targets can be expressed in different cell subsets, ensuring the therapy attacks all tumor cell subsets, or have opposite expression profiles in normal tissue, thereby decreasing on-target off-tumor effects and reducing toxicity. While IMMUNOTAR currently includes the features necessary for identifying such targets, a function could be incorporated to enable the systematic identification of dual target pairs, facilitating the development of dual therapies [55]. Further extensions of IMMUNOTAR could incorporate tumor single-cell RNA-sequencing profiling databases, such as the Single-cell Pediatric Cancer Atlas (scPCA) [56]. This would ensure the expression of dual targets across all tumor cell subtypes or confirm that at least one of the dual targets is expressed across the tumor [55].

While some tools focus on identifying tumor-specific proteins through alternative splicing and cancer-specific exons, IMMUNOTAR stands out for its ability to process both proteomics and RNA-sequencing data [57, 58]. Additionally, IMMUNOTAR integrates more public databases than existing tools, offering a comprehensive and quantitative evaluation to identify cancer-specific proteins. Overall, IMMUNOTAR fills a critical gap in the field by streamlining target identification, saving time and resources, and empowering researchers to focus on high-potential candidates for cancer immunotherapy development.

## Methods

### Enrichment databases included in IMMUNOTAR

#### Healthy tissue expression

An essential criterion for selecting a target is minimal to no expression in healthy tissues to mitigate on-target off-tumor effects. To quantitatively define this criterion, RNA-sequencing data from the Genotype-Tissue Expression (GTEx) database was incorporated [16]. The GTEx project conducted RNA-sequencing on 17,382 samples from 948 donors, encompassing 54 non-diseased tissue types [21]. The majority of GTEx samples are derived from donors aged 20 years and above. TPM-normalized tissue and sample-level RNA-sequencing data were retrieved from the GTEx portal. As GTEx is primarily comprised of adult tissue samples, RNA-sequencing data from pediatric tissues, obtained through the Evo-Devo Mammalian organs (Evo-Devo) project, was also included [17]. For both RNA-sequencing datasets, quantitative features were generated at both tissue and sample levels, resulting in a total of ten quantitative features per database. These features encompass the highest RNA expression value across tissues and individual samples, the count of tissue types and individual samples with expression > 10 TPM/RPKM, and a normalized maximum expression feature adjusted to the sample count annotated to a specific tissue type within each database. The threshold of 10 TPM/RPKM represents the 90th percentile cut-off, designating samples/tissues exceeding this value as high expressing for the protein. Given that certain immunotherapies do not penetrate the blood-brain barrier, we extracted summary statistics both with and without brain samples included. Acknowledging potential disparities between RNA expression and protein abundance, we incorporated proteomics data derived from 201 samples representing 32 distinct normal human tissues generated by Jiang et al. [18]. The processed protein quantification file at both the sample and tissue levels from the publication was queried. We extracted the same features from this database as the healthy tissue RNA-sequencing databases; however, the threshold for high-expression varied based on the 90^th^ percentile protein expression value.

#### Protein localization

While surface proteins play a crucial role as biological markers for diseases and therapeutic targets, their characterization is challenging due to their relatively low abundance, making it difficult to capture them using common sequencing or proteomics technologies [20]. To address this limitation, computational resources have emerged to aggregate and predict information, aiding researchers in defining protein localization. Among these databases, we have incorporated data from Compiled Interactive Resource for Extracellular and Surface Studies (CIRFESS) [20]. CIRFESS integrates various prediction strategies and annotations to assign a score to each protein based on the collected evidence supporting its classification as a surface protein, the confidence scores in CIRFESS range from 0-4. To broaden the list of surface proteins and leverage multiple protein localization databases, we also included data from the COMPARTMENTS database [19]. COMPARTMENTS utilizes available proteomics datasets and publications to assign scores to proteins based on the confidence that a protein is located on the plasma membrane, with confidence scores ranging from 0-5. In the IMMUNOTAR data matrix, the confidence score assigned to each protein, was included from both COMPARTMENTS and CIRFESS. Additionally, the length of the extracellular component of proteins was included as a feature in the IMMUNOTAR data matrix. This length information was obtained from the UniProt Knowledgebase, a database providing protein sequences along with functional details [59].

#### Biological significance

To gain deeper insights into the biological roles of protein targets in the queried cancer phenotype, gene dependency lists from the DepMap project were incorporated [22]. Within the DepMap project, a comprehensive genome-scale RNAi and CRISPR-Cas9 genetic perturbation screen was conducted to silence or knockout individual genes, identifying those genes that impact cell survival across 501 human cancer cell lines [22]. The DepMap database provides probability scores for genes in each of the surveyed cell lines. In the IMMUNOTAR data matrix, the quantitative feature included from DepMap is the probability score from the cell lines annotated to the cancer phenotype of interest. When there are multiple cell lines annotated to a single cancer phenotype, the user can decide if they want to average the probability across all cell lines or choose the maximum probability across all cell lines to populate the DepMap feature in the data matrix. Additionally, information from the Gene Ontology (GO) database was included to identify gene sets associated with pathways relevant to the cancer phenotype [23]. The quantitative feature for GO is binary, indicating whether the gene is present or not in the pathways of interest specified by the user.

#### Reagent/therapeutic availability

In the pursuit of identifying targets with established reagents specifically in pediatrics, the NCI relevant Pediatric Molecular Targets List (PMTL) was incorporated. This list aims to facilitate the development of safe and effective new drugs for pediatric cancer treatment (source: https://moleculartargets.ccdi.cancer.gov/mtp-pmtl-docs). The quantitative feature included for this database is binary, indicating whether the gene is present in the PMTL. To broaden the search for targets with developed reagents beyond pediatrics, the Therapeutic Target Database (TTD) and The Database of Antibody-drug Conjugates (ADCdb) were included. TTD provides information on drugs associated with diseases, the mode of action of developed drugs, and the stage of testing for these drugs [60]. ADCdb is a database dedicated to listing antibody-drug conjugates, a class of immunotherapeutic drugs commonly used in cancer, that are approved or are in development and their associated disease [25]. This information from TTD and ADCdb enables researchers to explore repurposing already developed drugs, if applicable. The feature extracted from both the databases is a score associated with the phase of development of the drug in the disease of interest. The defined scores for each development phase are listed in **Supplemental Table S12**.

### IMMUNOTAR feature extraction

#### User-input

In its initial phase, IMMUNOTAR constructs a data matrix with the initial columns being summary features of the user-input cancer expression dataset described in **Fig. 1B**. Using the input expression dataset, IMMUNOTAR can calculate the number of samples exhibiting “low-expression” of a gene. The user has the flexibility to either set their own threshold of “low-expression” or apply the default, below the 20th percentile of expression. Similarly, IMMUNOTAR can summarize the number of samples with “high-expression,” allowing the user to set a cut-off for “high-expression” or apply the default, above the 80th percentile of expression. Finally, IMMUNOTAR can provide a summary of the number of samples expressing the gene without any cut-offs for expression. This approach yields five features representing the cancer-expression data. While the default is to include all these features in the data matrix, users can choose to exclude some of these features. Additionally, users can introduce their own experimental data summary, such as performing a differential expression analysis between experimental groups. In this scenario, IMMUNOTAR utilizes all the columns in the user-provided dataset for the scoring matrix. Following the summarization of the cancer-expression dataset, the user has the option to rescale and handle missing values for the summarized data, employing methods detailed in **Fig. 1C**. If nothing is specified, the defaults for rescaling and missing values are applied to the dataset.

#### Enrichment databases

The subsequent step in the analysis focuses on enriching the scoring matrix with features from the public databases available within IMMUNOTAR. Users can choose specific databases for enrichment or default to enriching with all databases. For GO, DepMap, TTD, and ADCdb databases, additional user input for enrichment is required. To enrich with the GO dataset, users must list pathways of interest, allowing IMMUNOTAR to query related gene sets. For DepMap, TTD, and ADCdb enrichment, users need to feed in the cancer phenotype, enabling IMMUNOTAR to retrieve cancer-specific gene-dependency lists from the DepMap project and known targets for that cancer from TTD and ADCdb. Additional options for TTD and ADCdb enrichment include specifying the mode of action of the drug, limiting the search query to certain types of developed therapies. Mode-of-action categories can be found in **Supplemental Table S13**. Following enrichment, users can opt to rescale and fill in missing values for the enrichment features.

#### Applying curves and weights

To account for non-linearity of the feature values associated to scores, we included a curving factor that can be applied to each feature. The formula for curving the feature values is shown in **Fig. 1C**. This will allow the algorithm to better discriminate between values especially at the extreme ends of the scale and for the system to differentiate between subtle differences in the feature values. After applying curves, the user can also apply varying weights to each enrichment feature to highlight important features. The product sum of the curved feature value and weight for each gene is then calculated, generating the final gene prioritization score.

### IMMUNOTAR evaluation and optimization

#### MAP score

When extracting targets from TTD and ADCdb, some targets are associated with discontinued drug status and are given a negative score. Within IMMUNOTAR, targets that have a negative score are considered “known-negatives” and targets that have positive scores are considered “known-positives”. The MAP score for a phenotype that has both these types of targets is calculated as shown in **Fig. 2A**. The MAP score of the algorithm serves as a valuable metric for assessing the algorithm’s performance as well as providing an opportunity for optimizing the scoring parameters.

#### Optimization

The aim of this optimization is to identify features within the IMMUNOTAR scoring data matrix that characterize known-positive targets. While the optimization can only be applied to phenotypes with known or associated targets, once the parameters are generated, they can be applied to phenotypes lacking known or associated targets. We incorporate several mathematical methods for optimization, including Sequential Feature Selection (SFS), Genetic Algorithm (GA), Nelder-Mead (NM), and Brent’s Method (BM). SFS iteratively adds features to a model to improve its performance, effectively handling high-dimensional data by ensuring the inclusion of the most informative features. GA utilize principles of natural selection to explore large and complex search spaces, making them ideal for heuristic optimization [28]. NM algorithm, a simplex-based technique, is particularly useful for nonlinear optimization problems as it does not require gradient information [61]. Lastly, BM is a robust and efficient iterative technique for numerical root-finding, combining bisection, secant method, and inverse quadratic interpolation for faster convergence. The NM and BM methods are implemented using the optim library within R (https://www.R-project.org). The GA method was implemented using the GA R library [28].

### Datasets explored using IMMUNOTAR

#### Proteomic pediatric multi-cancer dataset and neuroblastoma (NBL) full proteome

Our overall application of IMMUNOTAR in our work is summarized in **Fig. 2C**. Initially, we applied IMMUNOTAR using a proteomics dataset that surveyed 949 cancer cell lines across 28 tissue types, encompassing approximately 40 histologically diverse cancer types [18]. This dataset serves as a comprehensive proteomics resource, offering the opportunity to investigate proteomic profiles across a wide spectrum of cancer phenotypes. Gonçalves *et. al.* conducted mass spectrometry, generating full proteome data for each cell line. To ensure result consistency, three sets of technical duplicates were run for each cell line. In total, 8,498 proteins were quantified across the dataset, with a median of 5,237 proteins quantified per cell line.

Given our specific focus on pediatric cancers, we refined the dataset using available meta-data to include only pediatric cancer cell lines that each had at least three representative cell lines. The processed data, averaged across technical replicates and provided through supplementary files in their publication, served as the input for IMMUNOTAR. For our neuroblastoma (NBL) specific analysis, the NBL full proteome data from this dataset was utilized. The NBL data surveyed 22 pediatric NBL cell-lines.

#### Pediatric Ewing sarcoma (EwS) surface proteomics

To validate the efficacy of our pipeline and optimized parameters and to compare our prioritization algorithm to ones used by other groups, we utilized a more focused surface proteomics dataset recently published by Mooney et al. [12]. This dataset was derived from tissue samples obtained from pediatric ES patient-derived or cell line-derived xenograft models [16]. A total of 19 samples were analyzed in technical or biological duplicates, depending on sample availability. Across all samples in the surface proteomics dataset, a total of 3,357 proteins were quantified. Within the publication, the team curated a list of 218 proteins that they annotated to surface proteins based on protein localization databases SurfaceGenie and UniProt. Next, they implemented a custom scoring methodology that combines data from their generated EwS specific surface and global proteome data, EwS RNA-sequencing data from the Pediatric Preclinical Testing Consortium (PPTC) and GTEx database to score the 218 proteins. This scoring methodology is not available as a tool to use for the research community. They identified novel potential candidates using this scoring mechanism that they proceeded to validate. For our methods, the processed proteomics data for each sample that was generated by the group and provided through the supplementary files of their publication was the input to IMMUNOTAR.

#### Multiple Myeloma (MM) surface proteomics

As a secondary analysis contributing to a cross-validation of our pipeline and optimized parameters in diverse experimental contexts and datasets, we applied our IMMUNOTAR pipeline to a surface proteomics dataset generated from MM cell lines by Ferguson et al. [11]. This dataset was derived from four multiple myeloma cell lines. The group ran technical duplicate or triplicate samples for each cell line for consistency of results. In total, 1,245 proteins were quantified in the surface proteomics cell line data. After quantification, the proteomics data was integrated with publicly available datasets including: RNA-sequencing data for MM cell lines from the Cancer Cell Line Encyclopedia (CCLE), RNA-sequencing data for blood cells from the Human Blood Atlas, and RNA-sequencing data for normal tissue from GTEx. For surface protein localization they included the Compartments database. They extracted 5 quantitative features from these databases related to surface abundance and specificity for plasma cells to identify immunotherapeutic targets for MM [11]. For their scoring algorithm, they included all genes identified in the surface proteomics data and additionally genes identified in MM bulk RNA-sequencing data, thus evaluating around 33,000 candidates. Using their scoring algorithm, they identified novel MM specific therapeutic candidates that they moved forward to validate in the lab. For our methods, we utilized the quantified proteomics files from the supplementary data and ran it through IMMUNOTAR.

## Data Availability

Data utilized in this study are publicly available and can be accessed as follows: The pan-cancer proteomics map protein intensity data and meta data were downloaded from cellmodelpassports.sanger.ac.uk as referenced by the publication (PMID:35839778) [26]. The EwS surface proteomics data was downloaded from Supplementary Data S1 table and the scoring metric for this dataset was downloaded from Supplementary Data S3 (PMID: 37812652) [12]. The Multiple Myeloma surface proteomic data was downloaded from Supplementary Data 1 and the scoring metric for this dataset was downloaded from Supplementary Data 2 (PMID: 35840578) [11].

## Code Availability

IMMUNOTAR code is freely available on GitHub: https://github.com/sacanlab/immunotar

## Acknowledgements

This work was supported by National Institutes of Health (NIH) grants U54-CA232568 (JMM), R01-CA237562 (SJD), R35-CA220500 (JMM), and T32-CA009140 (AKW). This work was delivered, in part, by the NexTGen Cancer Grand Challenges partnership funded by Cancer Research UK (CGCATF-2021/100002) and the National Cancer Institute (CA278687-01) and The Mark Foundation for Cancer Research. B. Mooney is funded by a trainee award from the Michael Smith Health Research BC (RT-2023–3194).

## Author Contributions

Conceptualization: RS, SJD, JMM, AS; Methodology: RS, KLC, BM, AKW, SJD, JMM, AS; Formal analysis: RS; Investigation: RS, KLC, BM, JMM, SJD, AS; Validation: RS, BM; Writing – Original Draft: RS; Writing – Review and Editing: KLC, BM, AKW, JMM, SJD, AS; Supervision: SJD, JMM, AS; Funding Acquisition: JMM, SJD.

