## Supplemental Figures S1-S2 for "IMMUNOTAR - Integrative prioritization of cell surface targets for cancer immunotherapy"

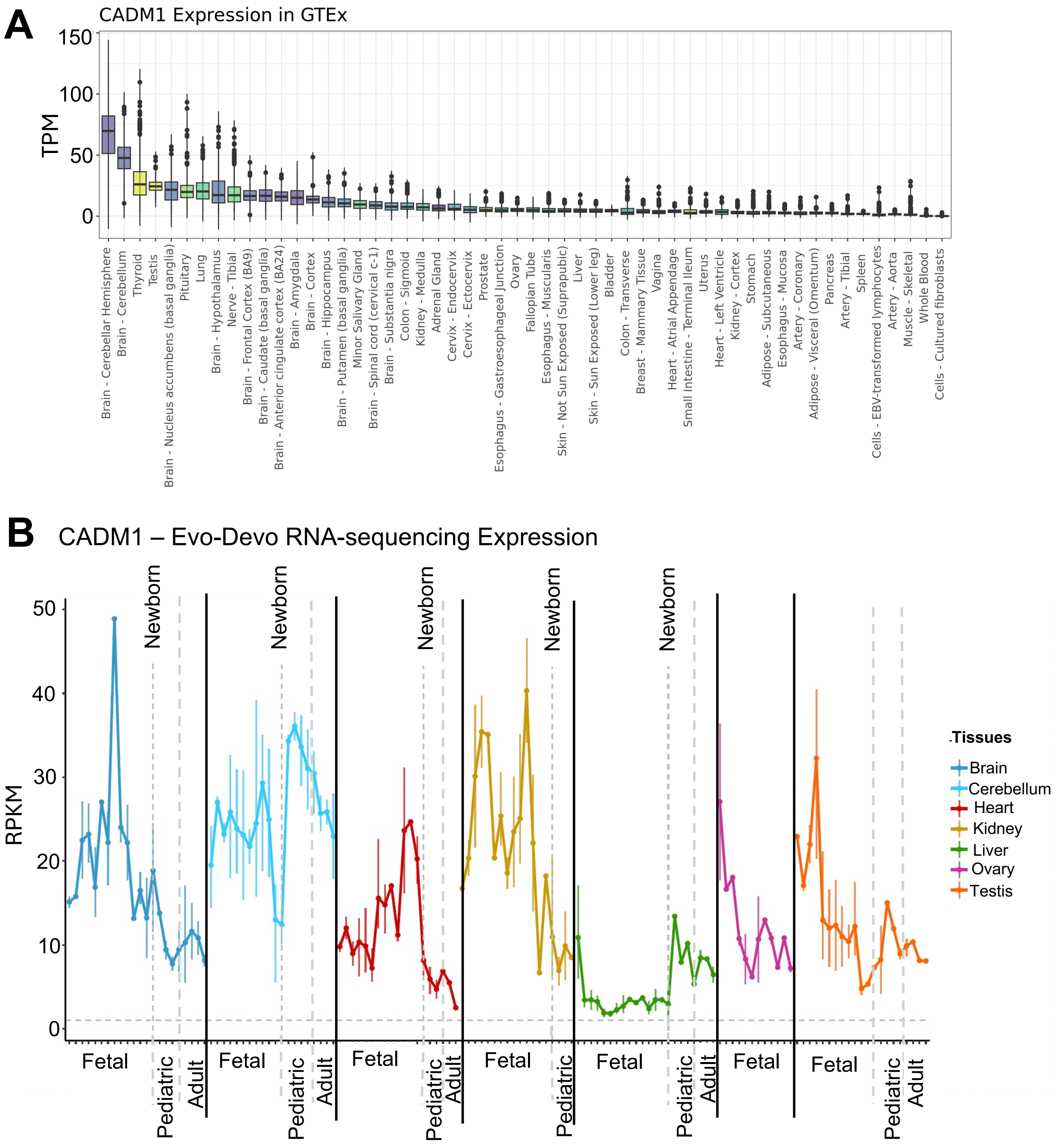


**Supplemental Fig. S1: *CADM1* expression in normal tissues databases GTEx and Evo-Devo.** **A)** Normal tissue expression per GTEx RNA-sequencing database. **B)** Normal tissue expression per Evo-Devo RNA-sequencing database.

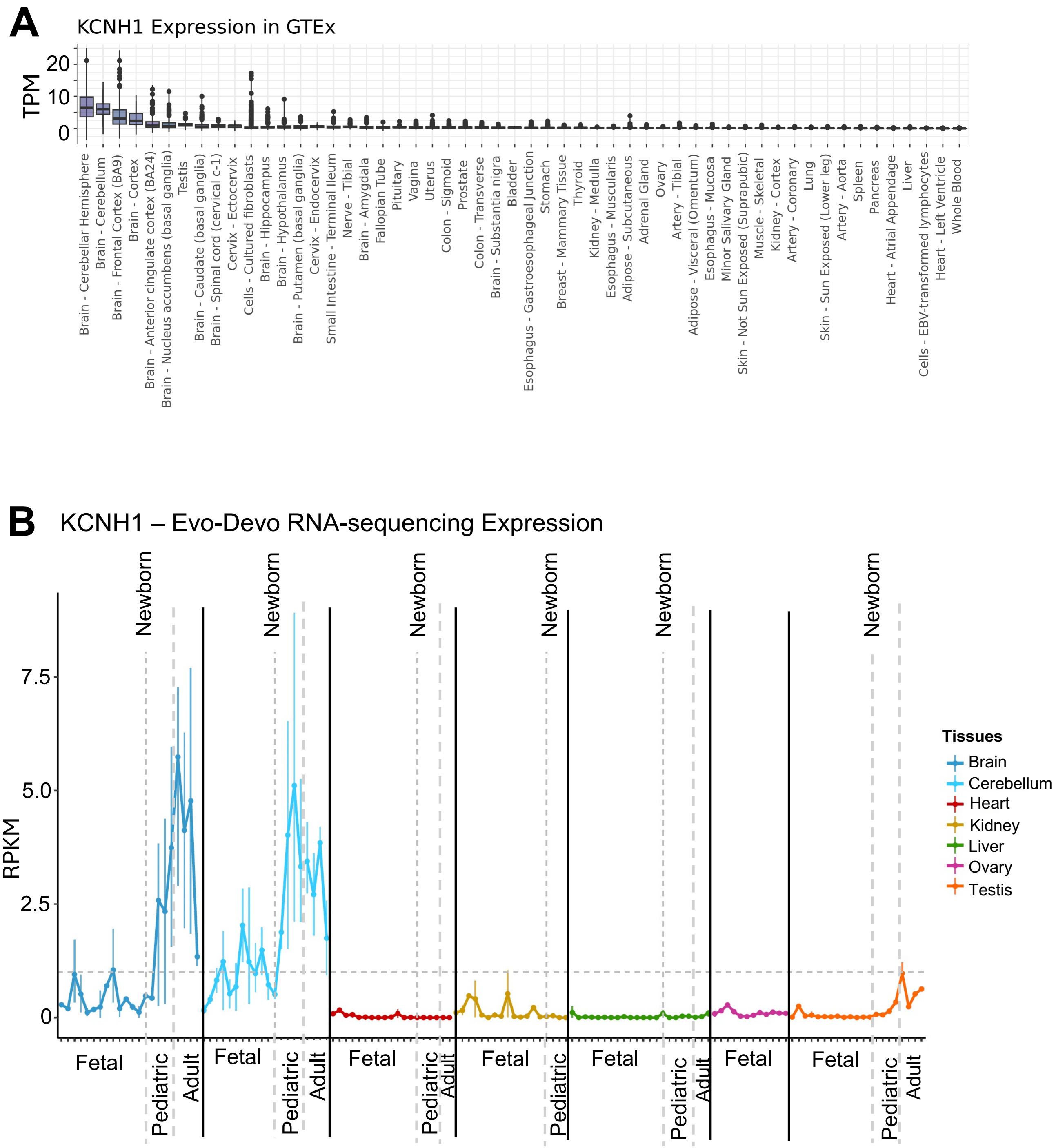


**Supplementary Fig. S2**: ***KCNH1* expression in normal tissues databases GTEx and Evo-Devo**. **A)** Normal tissue expression per GTEx RNA-sequencing database. **B)** Normal tissue expression per Evo-Devo RNA-sequencing database.
