## Supplemental File S2 for "IMMUNOTAR - Integrative prioritization of cell surface targets for cancer immunotherapy"

#Supplemental File S2: IMMUNOTAR default analysis parameters values
##Please refer to Supplemental Table S2 for each feature description

handlenan: '0ormin'   #change default handlenan. (if not given in a project configuration, the code defaults to 'none'). If handlenan=0 and a column contains negative values, the code will print a warning. If handlenan='0ormin', we use the min value if there are any negative values or 0 if everything is positive.
colhandlenan: #The user can specify handlenan for individual columns
  healthyprot: 'min'

rescale: 'percentile' #This will rescale all feature values 
colrescale: #Rescaling specific columns 
  cirfess: 'rangemap0100'
  compartments: 'rangemap0100'
  depmap: 'rangemap0100'
  opentargetsurface: 'rangemap0100'
  theratarsurface: 'rangemap0100'

enrich: #parameters to be passed into data_enrich()
#only passing in databases that don't need additional parameters to enrich with 
  enrichtypes: 'gtex,evodevo_pediatric,healthyprot,compartments_sp,cirfess_spc,uniprot,opentargetsurface'  #by default, we use data_enrich. If you do not want to use enrich, define doenrich=0 as a top level attribute, or define enrich.enrichtypes=''
  #some enrich types require additional arguments. provide them here. examples are below but commented out. 
  #depmapids: 'Rhabdomyosarcoma' #this can be numeric id(s) or text(s) to search.
  #goids: 'Notch binding'
  #go_collate: 'merge' #separate|merge determines whether we create separate columns for each goid.

colweight: #specify weight for individual columns
  expr: "+1 each" #Will apply a +1 weight to each feature extracted from cancer expression dataset 
  expr_numlowexpressed: -1
  gtex: "-1 each" #Will apply a -1 weight to each feature extracted from GTEx 
  evodevo_pediatric: "-1 each" #Will apply a -1 weight to each feature extracted from Evo-devo
  healthyprot: "-1 each" #Will apply a -1 weight to each feature extracted from Healthy proteomics dataset  
  compartments_sp: +1
  cirfess_spc: +1
  uniprot: +1
  opentargetsurface: +1
  theratarsurface: +1
  depmap: +1

curve: 0 #No curving is applied in the default analysis 
