## Supplemental File S3 for "IMMUNOTAR - Integrative prioritization of cell surface targets for cancer immunotherapy"

# Configuration may contain a single 'dataset' entry, or a 'datasets' containing a list of them.
# A dataset can contain these attributes (and any option that data_summarize() function accepts):
# datafile, summarytype.
# The datafile may contain {datadir} are {userdatadir}, which are replaced with myconfig('datadir'), etc.
# File names (unless they already exist in the cwd) are adjusted relative to the folder containing this yml file.
# All parameter values will be imported from the import files listed below unless changed in this file. 

importdefaultparams: T  #this is equivalent to using importfiles to import the default params (demonstrated below).
importoptimizedparams: T #this is equivalent to using importfiles to import the optimized params (demonstrated below).

#You can import other configuration files using "importfiles" entry. See the content of the imported files for the options you can change.
#The files listed below can alternatively be imported by setting importdefaultsparams=T and importoptimizedparams=T. Re-listed here for demonstration.
importfiles:
 - 'project_defaultparams.yml'   
 - 'project_optimizedparams.yml'

# A dataset is your experimental data that forms the starting point for an analysis project.
# Experimental data can be specified using "dataset" (single dataset) or "datasets" (multiple datasets) entry.
# If you want to only use the enrichment databases (without any experimental datasets), create a dataset 
# with gene names and without any numerical data. Alternatively, use dataset: '__ALLHUMANGENES__' or dataset: '__ALLHUMANSURFACEGENES__'.

#dataset: '__ALLHUMANGENES__'
dataset: '__ALLHUMANSURFACEGENES__'

enrich: #parameters to be passed into data_enrich(). For now these parameters are being fed from the default parameters file (Supplemental File S2)
  #some enrich types require additional arguments. provide them here.
  depmapids: 'Rhabdomyosarcoma' #this can be numeric id(s) or text(s) to search
  #goids: 'Notch binding'
  #go_collate: 'merge' #separate|merge determines whether we create separate columns for each goid.


rescale: 'percentile' #change default scale. (default is 'percentile')
#colscale: #specify scale for individual columns
 # gtex: 'percentile' #linearly scales min..max to 0..100
  #go: 'none'

weight: 1 #Set the default weight. If you set this, remember to override this default value for columns that should have an opposite signed weight.
  
#coloptions: #Options may also be organized for each column. These will only be effective if the same option is not specified in one of the colXXX option groups (e.g., colscale, colweight, etc.)
  #gtex:
    #scale: 'percentile' #if both colscale.gtex and coloptions.gtex.scale are provided, colscale.gtex is used and coloptions.gtex.scale is ignored.
    #weight: 0.5


# If there are certain genes that you consider as known targets, list them here. For these genes, we
# calculate the MAP (mean average precision) score indicating how well the analysis was able to detect them as targets.
# Provide a list or a CSV text.
knownpositives: 'ALK','MEGF10','GPC2'

#After the analysis, if you would like extract a table of percentile_rank's for certain genes, use 
# Provide a list or a CSV text.
reportforgenes: 'ALK,GPC2'

#After the analysis, if you would like to write the results into a file, provide the filename here.
#outfile: 'demo_project_result.xlsx'
